# The interplay between small RNA pathways shapes chromatin landscapes in *C. elegans*

**DOI:** 10.1101/320713

**Authors:** Ekaterina Gushchanskaia, Ruben Esse, Qicheng Ma, Nelson Lau, Alla Grishok

## Abstract

The nematode *C. elegans* contains several types of endogenous small interfering RNAs (endo-siRNAs) produced by RNA-dependent RNA polymerase (RdRP) complexes. Both “silencing” siRNAs bound by Worm-specific Argonautes (WAGO) and “activating” siRNAs bound by the CSR-1 Argonaute require the DRH-3 helicase, an RdRP component. Here we show that, in the *drh-3(ne4253)* mutant deficient in RdRP-produced secondary endo-siRNAs, the silencing histone mark H3K9me3 is largely depleted, whereas in the *csr-1* partial loss-of-function mutant this mark is ectopically deposited on CSR-1 target genes. Moreover, we observe ectopic H3K9me3 at enhancer elements in both *drh-3* and *csr-1* partial loss-of-function mutants and describe small RNAs matching enhancers. Finally, we detect accumulation of H3K27me3 at highly expressed genes in the *drh-3(ne4253)* mutant, which correlates with their reduced transcription. Our study shows that when abundant RdRP-produced siRNAs are depleted, there is ectopic elevation of noncoding RNAs linked to increase in silencing chromatin marks. Moreover, our results suggest that enhancer small RNAs may guide local H3K9 methylation.

## INTRODUCTION

The RNA interference machinery, and specifically Argonaute proteins that bind small RNAs, are present both in the cytoplasm and the nucleus in diverse species (1,2). In fission yeast and plants, there is an elaborate connection between siRNAs and heterochromatin formation (3–5). In animals, Piwi-interacting RNAs (piRNAs) that protect the germline from parasitic elements are strongly implicated in triggering chromatin-based silencing (1). In the nematode *C. elegans,* as well as in mammalian cultured cells, exogenous or transgene-based introduction of double-stranded RNA (dsRNA) can initiate transcriptional silencing and/or heterochromatin formation (6–9). Finally, there are endogenous siRNAs (endo-siRNAs) produced from the dsRNA segments arising from large hairpins, overlapping 5’ or 3’ ends of transcripts, or repetitive elements that have been detected in oocytes of higher animals (10,11). Whether they also regulate chromatin compaction is an open question.

Open chromatin conformation at enhancer elements is an essential requirement for enhancers to regulate global and tissue-specific transcription, and is conserved across species, including *C. elegans* (12). Enhancers constitute stretches of DNA with characteristic features of actively transcribed chromatin, such as DNase1 hypersensitivity, high occupancy by transcriptional machinery, and specific histone modifications (H3K4me1, H3K27ac) (13–15). Intragenic enhancers promote local gene transcription, whereas intergenic enhancers can undergo chromatin looping to promote distal gene transcription, up to several thousand base pairs away. Beyond their initial discovery (16–18), new features of enhancers have emerged that require further dissection to fully understand enhancer function.

For example, the field now appreciates that bidirectional transcription at enhancer regions (19–21) results in production of noncoding RNAs (ncRNAs), named enhancer RNAs (eRNAs). This transcription appears to play an important role in enhancer function, most notably by keeping chromatin open (13,15), although specific examples of the functional significance of eRNA sequences themselves are also known (15). The most prominent examples of enhancer-derived ncRNAs are RNAs produced from the *Ultrabithorax* domain in the homeobox gene region in *Drosophila* (22) and the Evf-2 ncRNA associated with the Dlx-5/6 ultraconserved enhancer in mouse (23). Finally, super-enhancers are spatially and functionally associated with miRNA genes (24).

There is a potential for nuclear siRNA production from the dsRNA arising from bidirectional transcription at enhancers (15). If such siRNAs bind Argonaute proteins guiding heterochromatin marks, then this mechanism may serve a regulatory purpose to make enhancers closed in some cell types, or this may simply be an unwanted aberration. In either case, such siRNAs need to be controlled. Putative enhancer elements have only recently been annotated in the *C. elegans* genome (25,26), and here we provide first evidence of small RNAs matching enhancers.

The most abundant *C. elegans* endo-siRNAs are 22G-RNAs produced through amplification of primary siRNAs by RNA-dependent RNA Polymerases (RdRP) (27). A group of redundant Worm-specific AGO proteins (WAGO) bind 22G-RNAs and target repetitive elements, pseudogenes, transposons, and some duplicated protein-coding genes for silencing (27). Two nuclear WAGOs are implicated in facilitating transcriptional silencing: NRDE-3 in the soma (28), and HRDE-1 in the germline (29). Both are capable of inducing H3K9me3 at genes complementary to exogenous dsRNA or at transgenes downstream of piRNAs (6,8,28–33). However, the interconnection between 22G-RNA production, H3K9 methylation and transcriptional silencing at the endogenous WAGO targets remains unclear (34,35).

Whereas the concept that nuclear Argonaute proteins guide silencing histone modifications is well established, the CSR-1 (Chromosome Segregation and RNAi deficient) Argonaute (36,37) does not fit this stereotype. It is present both in the cytoplasm and the nucleus, it binds 22G-RNAs that are largely antisense to highly and widely expressed genes, and it is associated with nascent transcripts and mature mRNAs complementary to these siRNAs. In contrast to direct gene-silencing roles for WAGOs, CSR-1 has an “activating” or “anti-silencing” role, since nascent transcripts complimentary to CSR-1-bound siRNAs decrease in *csr-1*-deficient worms (38). Previously, we showed by Global Run-on Sequencing (GRO-seq) that in *csr-1* and *drh-3* partial loss-of-function mutants there is both a decrease in nascent transcripts targeted by CSR-1 22G-RNAs and a genome-wide increase in antisense transcription (38). Notably, DRH-3 (Dicer-related helicase) is a component of RdRP complexes producing 22G-RNAs and is required for generation of both WAGO-bound and CSR-1-bound small RNAs (27).

Here, we report that a decrease in transcription of CSR-1 target genes observed earlier (38) correlates with ectopic silencing chromatin marks. Moreover, we describe an increase in noncoding transcription and elevation of H3K9me3 at putative gene enhancers in both *csr-1* and *drh-3* mutants. Finally, we identify small RNAs originating from enhancers in previously published datasets. Our results support a model whereby “activating” and “silencing” secondary siRNAs in *C. elegans* have complementary functions required for proper genome-wide chromatin organization. Our data also suggest that, in the absence of abundant 22G-RNAs, rare small RNAs, such as those produced from enhancers, may be incorporated into Argonaute complexes capable of guiding H3K9me3 deposition.

## MATERIALS AND METHODS

### Strains

Strains were maintained at 20°C unless otherwise noted, using standard methods (39). Bristol N2 was the WT strain used. Mutants used in this study were *drh-3(ne4253),* and partially rescued *csr-1(tm892*) (strain WM193). For the ChIP-seq experiments, worm populations were synchronized and grown for approximately 40 h after hatching at 20°C on OP-50 *E. coli* at a density of ~50,000 animals per 15-cm Petri dish until they reached L3 stage.

### Chromatin Immunoprecipitation

Chromatin immunoprecipitation was performed following the modENCODE Protocol from the Lieb Lab (40) with some modifications. The worm pellet fixed with 2% PFA for 30 minutes at 20°C was washed in M9 3 times and resuspended in RIPA buffer supplemented with Protease inhibitors (Thermo Scientific, 78443). DNA fragmentation was performed using sonication with Covaris (Peak power 240, Duty factor 20, Cycles/burst 200, 8 min). Then, 1.5–2mg of cross-linked chromatin extract was incubated at 4°C ON with a specific antibody and the immune complexes were then incubated with 50µl IgG Dynabeads (Invitrogen) for 3h at 4°C. DNA was cleaned up with the Qiagen PCR purification kit. We used the following antibodies: 5μg of anti-H3K27me3 (Diagenode, pAb-195-050), 5μg of anti-H3K36me3 (Abcam, ab9050), 5μg of anti-H3K9me3 (Abcam, ab8898). Validation information for the commercial antibodies is included at the manufacturers’ websites.

### Library preparation, sequencing and data processing

ChIP-seq libraries were prepared using TruSeq Illumina kit (set A – 15034288, set B – 15034289) according to manufacturer’s instructions. Sequencing was performed on Illumina NextSeq500. The 75-bp single-end Illumina sequencing reads were preprocessed by trimming the adapter sequences with Cutadapt (41). After that, reads were aligned to the WS220/ce10 assembly of the *C. elegans* genome using Bowtie for Illumina (Galaxy Version 1.1.2) (42) with default settings. The SAMtools (Galaxy Version 1.1.2) (43,44) utility was used to convert the alignments to BAM format. A table containing the number of reads aligned to genome (Supplementary Table 1) demonstrates sufficient sequencing depth (~14-28M of reads) Aligned reads were visualized in IGV browser (45,46) to check the overall read distribution shape. Although not a quantitative metric, visible enrichment at known binding regions confirms successful ChIP-seq experiment.

Two independent ChIP experiments were performed with each type of antibody. To estimate the concordance of 2 replicates, we applied Pearson’s correlation coefficient demonstrating perfect linear correlation (close to 1) between most replicates and good correlation (>=0.68) between H3K9me3 wild type replicates and H3K27me3 *drh-3(ne4253)* replicates (Supplementary Table 2). Duplicates reads were removed, and the data were processed as described below. Sequencing data has been submitted to GEO database (submission Series GSE115629).

### Analysis of ChIP-seq data

Aligned ChIP-seq data was normalized TMM normalized using edgeR package. Read counting at specific loci (either genes, enhancers or non-coding genomic regions) was performed with package GenomicAlignments (45); only reads with mapping quality 20 or higher were included in subsequent analyses. Obtained coverage was expressed as the log2-transformed ratio of the RPKM value in the immunoprecipitated DNA sample after subtraction of the RPKM value in the non-immunoprecipitated (input) DNA. Only regions in which the normalized count value in the immunoprecipitated DNA sample was higher than that in the corresponding input DNA in at least one sample in the set were considered. Gene identifiers were converted to Refseq mRNA IDs using the WormBase Converter utility (http://wormbasemanager.sourceforge.net/) and the DAVID Gene ID Conversion Tool (47).

### Analysis of GRO-seq data

GRO-seq reads (38) were aligned to the WS220/ce10 assembly of the *C. elegans* genome using Bowtie for Illumina (Galaxy Version 1.1.2) (42) with default settings. Reads matching ribosomal RNA loci were removed, as described before (48). Read counting at genomic regions was performed with package GenomicAlignments (49); only reads with mapping quality 20 or higher were included in subsequent analyses. Regions without reads across the sample set were removed. Counts were then normalized using the TMM method, which takes RNA composition bias into account (50), using the edgeR package (51). Coverage was expressed as RPKM (reads per kilobase per million mapped) and log2-transformed.

### Heatmaps

The purpose of heatmaps is the visualization of changes in histone modification along the gene-coding region in mutants compared to wild type. To generate the heatmaps, Galaxy platform was used. The aligned reads were normalized to RPKM using bamCoverage option (bin size=100bp, scaling/normalization method=rpkm, no scaling factor, ignoring missing data, reads extension=250, ignore duplicates=true, center regions with respect to the fragment length=false, minimum mapping quality=1) and then normalized to input using bigWigCompare option (method of comparison=ratio, pseudocount=1.0). Further, mutant coverage representing one of the histone modifications was compared with wild type coverage for corresponding modification and belonging to the same replicate (method of comparison=log2, pseudocount=1.0). The mean of two replicates for either H3K27me3 or H3K36me3 was then visualized along the gene bodies using computeMatrix (Distance in bases to which all regions are going to be fit=1000 bp, Distance upstream/downstream of the region start position=+/-200, Length, in bases, of non-overlapping bins used for averaging the score over the regions length=50bp, Convert missing values to 0=true) option and plotHeatmap (Sort regions=no ordering, Color map to use for the heatmap =bwr, Heatmap width in cm=10.0, Heatmap height in cm =20.0, What to show=heatmap and color bar). The options were standard if not specified otherwise. Gene subsets used for the analysis are listed in the Supplementary Table 3.

### Analysis of putative enhancer regions and ATAC-seq peaks

Putative enhancer regions were obtained by combining coordinates of regions from domains 8, 9 and 10 (26). Enhancer domains at least 1500bp distal to any annotated transcription start site or transcription termination site were considered distal enhancer domains. Enhancer domains intersecting coordinates of genes < 15kb by at least 50bp were considered intragenic enhancer domains. Among 177845 regions detected in (25), 13602 were considered putative enhancers. After filtering, percentage of enhancer with positive value of H3K9me3 coverage after input subtraction at least in one sample was 56%.

ATAC-seq peak coordinates (25) were downloaded from the NCBI GEO database (GSE89608). Distal and intragenic ATAC-seq peaks were obtained as for enhancer domains. Among 56624 total genomic regions, 9448 ATAC-seq enhancer regions were defined. After filtering, positive values of H3K9me3 coverage after input subtraction at least in one sample were detected for 46% of ATAC-seq peaks. The portion of the genome not containing ATAC-seq peaks was divided into genome regions of 2.5kb.

### Small RNA bioinformatics analyses

Small RNA library sequences were downloaded from the NCBI Gene Expression Omnibus and Sequence Read Archive under the following accessions: GSM454002, GSM455389, GSM455390, GSM455387, GSM455388, SRR030720, SRR030721, SRR030722, SRR030719, and SRR030717. Fastq files were validated and adapters were trimmed with the programs Fastqc (52) and Cutadapt (41), respectively. The genome and Refseq transcript assemblies for “ce10” were loaded into our custom “gene-centric process” small RNA analyses pipeline (53,54). This program maps small RNAs with Bowtie1 (42) with a maximum of 2 base mismatches, counts the genomic strand reads against the Ensembl gene transcript models using Bedtools (55) using the longest transcript isoform as the main gene model, and generates WIG plots for coverage visualization on the UCSC Genome Browser (56). Each library’s read counts were normalized against library depths (counts per million, cpm), as well as frequency of genomic locations (cpm / number of genomic loci).

For further analysis small RNA read coordinates were intersected with enhancer domain coordinates (26). For graphical representation purposes, the total number of reads for each sample was reduced to the number of reads observed in the sample with minimal read coverage (i.e. down sampling was performed). Percentages of reads were obtained and grouped in blocks with each block representing 1500 reads.

## RESULTS

### Competing regulation of H3K9me3 on highly active genes by the CSR-1 and WAGO pathways

Previously, we observed similar alterations in global transcription in the viable partial loss-of-function *csr-1* and *drh-3* mutant larvae compared with wild type animals (1). To establish a connection between transcription changes and chromatin status in these mutants, we performed ChIP-seq with antibodies specific to several histone modifications: H3K27me3, H3K36me3, and H3K9me3.

Consistent with the published data (57), H3K9me3 was found predominantly at chromosome arms (Figure 1A). Whereas the level of H3K9me3 increased at CSR-1 target genes in the *csr-1* mutant compared to wild type (Figures 1B, left plot, 1D, and S1), in *drh-3(ne4253)* we observed a depletion of this silencing mark at the same genes (Figure 1B, right plot). Because both the CSR-1 and the WAGO pathways are deficient in *drh-3(ne4253)*, these results suggest a competition between the “activating” and “silencing” secondary siRNAs, such that the WAGO pathway promotes H3K9me3 and CSR-1 inhibits its deposition at the majority of CSR-1 target genes. This type of competitive relationship has been suggested by other studies utilizing transgenic strains (30,33,58). Consistently, we observed a reduction in H3K9me3 modification abundance in d*rh-3(ne4253)* by western immunoblotting, which indicates that 22G-RNA-dependent methylation of H3K9 is globally affected in this mutant. On the contrary, in the *csr-1* mutant, there was a slight increase in H3K9me3 detected by western, suggesting that the WAGO silencing pathway is overactive under these conditions (Figure 1C, leftmost blot). We were not able to test whether this increase in H3K9me3 is dependent on DRH-3, because combining *drh-3(ne4253)* with partially rescued *csr-1(tm892)* mutant (strain WM193, (36)) led to early developmental arrest. In the meantime, we have not observed global changes in the levels of H3K27me3 and H3K36me3 in the mutants compared to wild type as assessed by Western blotting (Figure 1C, middle and rightmost blots).

**Figure 1.**
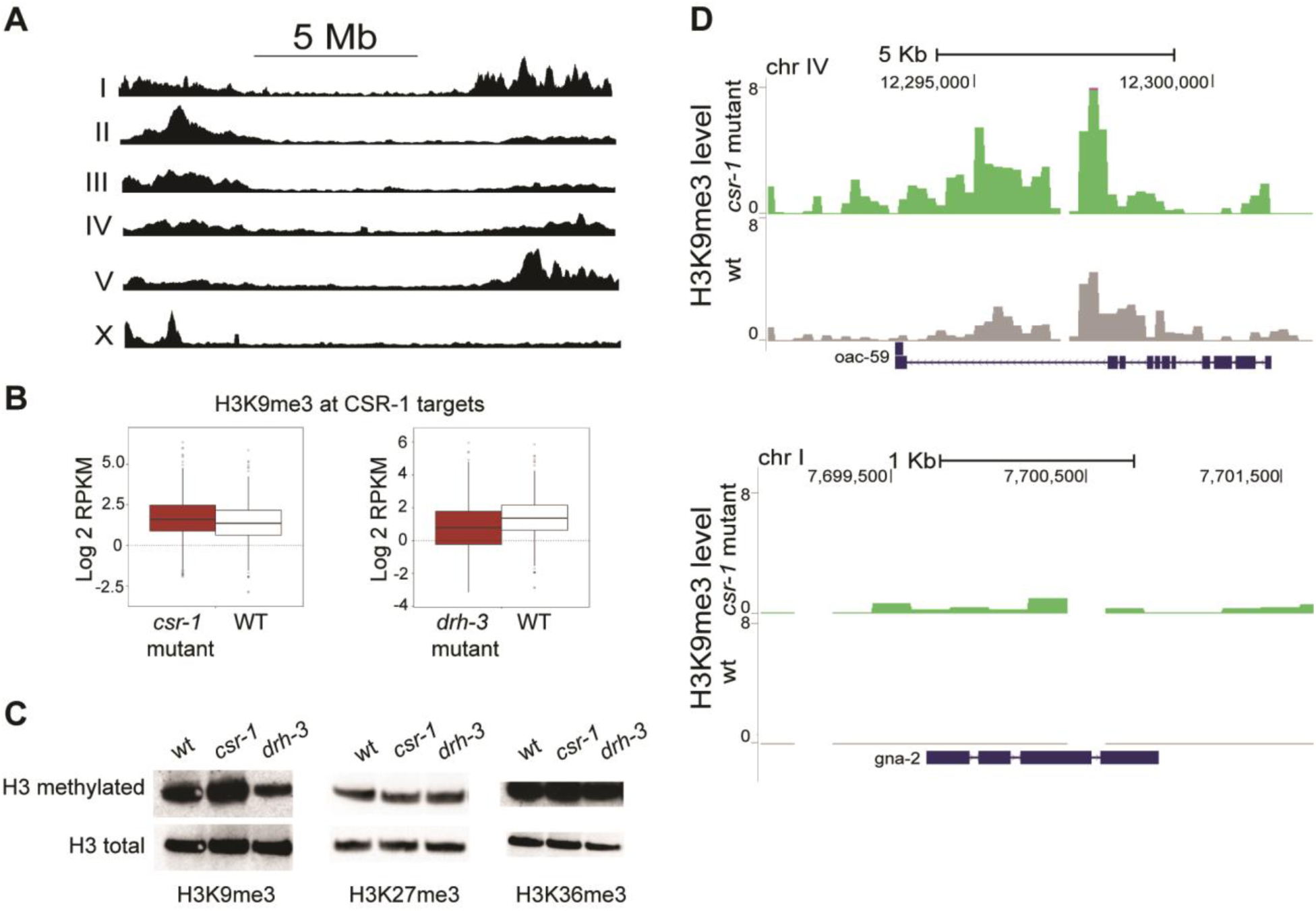
H3K9me3 alterations at CSR-1 target genes in partially rescued *csr-1(tm892)* and *drh-3(ne4253)*. **(A)** On the *C. elegans* autosomes, H3K9me3 is enriched at the chromosome arms and, on the X chromosome, only at the left arm. Read coverage in wild type worms (L3) normalized to, sequentially, RPKM and input is represented. **(B)** Box plot demonstrating an increase of H3K9 methylation at the CSR-1 target genes in *csr-1* mutant, but not in *drh-3 (ne4253)*. ChIP-seq data were normalized as described in methods. P-values<2.2*10^−16^(Wilcoxon test). **(C)** Global level of H3K9me3 in L3 stage *csr-1* mutant increases compared to wild type. On the contrary, H3K9me3 level in *drh-3* mutant is reduced. The levels of H3K27me3 and H3K36me3 are not changing globally. The upper blots show H3K9me3/H3K27me3/H3K36me3 histone modifications detected with following antibodies: ab8898 (Abcam), 07-449 (Millipore) and ab9050(Abcam), respectively. For control, the membrane was stripped and re-blotted with anti-H3 antibody (Abcam, ab1791). **(D)** Coverage tracks demonstrating two CSR-1 target genes of different sizes and exon/intron composition. In wild type (grey), H3K9me3 is very low (upper plot) or not detected (lower plot), whereas in *csr-1(tm892)* mutant (green), an increase in H3K9me3 is observed. TMM and CPM normalization was performed followed by input reads subtraction from each sample.

Our results are consistent with the idea that, in the *csr-1* mutant, WAGO-induced H3K9me3 is likely to play a role in inhibiting active genes (59,60). Therefore, CSR-1-bound siRNAs may protect their complementary pre-mRNAs from the WAGO-dependent 22G-RNAs guiding H3K9 methylation (Figure 6).

### Increased ncRNA production in *csr-1* and *drh-3* loss-of-function mutants correlates with accumulation of silencing marks at enhancers

Since there is a global increase in antisense transcription detected by GRO-seq in the *csr-1* and *drh-3* loss-of-function mutants (38), we chose to examine a specific CSR-1 target gene, cyclin E (*cye-1*), as antisense transcription has been previously detected at this locus (61). In agreement with the genome-wide analyses (38), we observed a decrease in the number of reads corresponding to sense transcription of the *cye-1* gene in *csr-1* and *drh-3* mutants (Figure 2). We also observed an increase in antisense reads corresponding to the large intron 2 of *cye-1* (Figure 2). Notably, intronic enhancer elements have been recently annotated in *C. elegans* (26), including the second intron of *cye-1*. Therefore, *cye-1* antisense transcript may represent an enhancer RNA.

**Figure 2.**
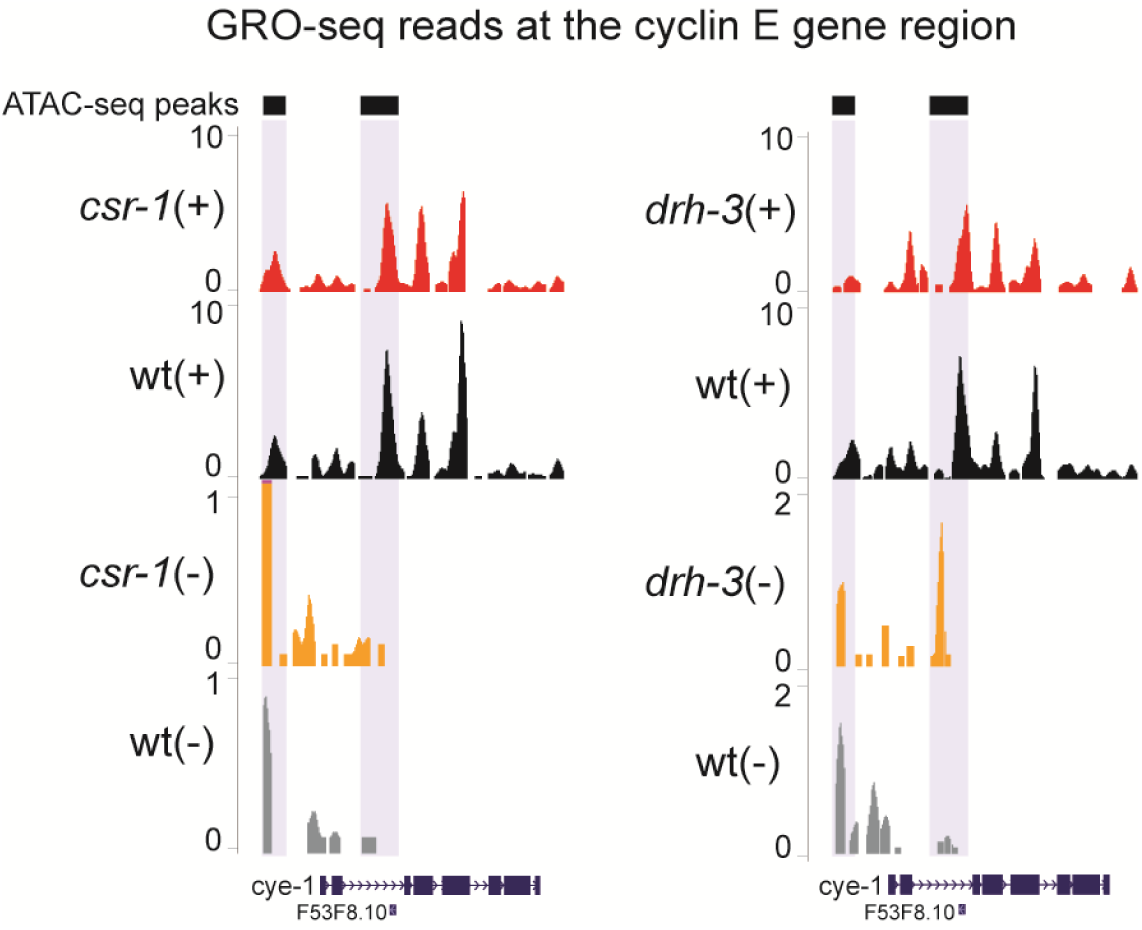
GRO-seq reads distribution at the *cye-1* gene region. A reduction in GRO-seq reads number on the plus strand in *csr-1* mutant *(left plot)* and *drh-3* mutant (*right plot*) compared to wild type (red vs. black) corresponds to the directionality of *cye-1* gene transcription. On the minus strand, at the intronic enhancer region detected by (25), there is an increase in GRO-seq reads in *csr-1 (left plot)* and *drh-3* (*right plot*) mutants compared to wild type (orange vs. grey). Normalization was performed as described in methods.

To further analyze transcription associated with other potential enhancer regions, we re-analyzed enhancer-mapping data from two recent publications (25,26). In (25), open chromatin regions associated with enhancers were detected by ATAC-seq and the functionality of some was confirmed. In another study (26), enhancer locations were predicted using established histone modification signatures. Our analysis of GRO-seq data identified a global increase in transcription at ATAC-seq-mapped distal (intergenic) enhancers in both *csr-1* and *drh-3* mutants (Figure 3B). We obtained similar results having analyzed GRO-seq reads at predicted enhancers (26) (Figure 3C). Indeed, we found increased antisense transcription at a functional upstream regulatory region of the *daf-12* gene that was more recently defined as active L3 stage enhancer by ATAC-seq (25,62) (Figure 3D).

**Figure 3.**
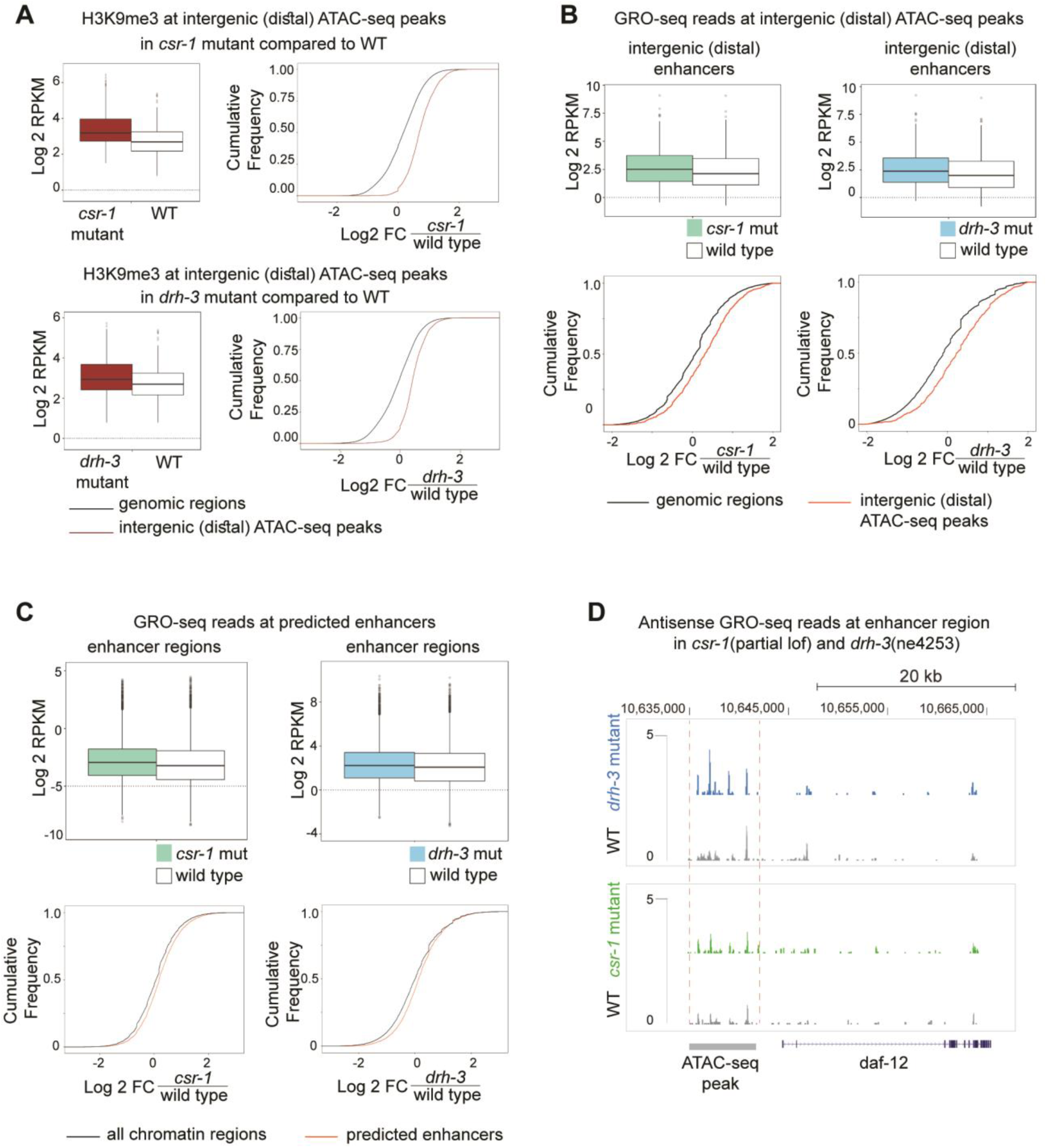
Elevated H3K9me3 and transcription at enhancer regions in partially rescued *csr-1(tm892)* and *drh-3(ne4253)* mutants. **(A)** Box plots and cumulative plots demonstrating H3K9me3 increase at distal (intergenic) enhancers detected by ATAC-seq (25) in *csr-1* and *drh-3* mutants compared to wild type. ChIP-seq data normalization was performed as described in methods. The genome-wide regions not containing ATAC-seq peaks were defined by dividing the genome into windows (genomic regions) of 2.5 Kb. P-values<2.2*10^−16^(Wilcoxon test). **(B)** GRO-seq read coverage is increased at distal enhancer regions (intergenic ATAC-seq peaks) in *csr-1* (left panel) and *drh-3* (right panel) mutants. P-value=8.694*10^−13^ and P-value<2.2*10^−16^ are corresponding to the left and the right plot respectively (Wilcoxon test). GRO-seq data from (38) were normalized as described in methods, and log2 transformed RPKM was plotted. P-values <2.2*10^−16^ (Wilcoxon test). Cumulative plots confirming the GRO-seq data analysis for *csr-1* (on the left) and *drh-3* (on the right) mutants and demonstrating an increase in GRO-seq reads coverage in both mutants at distal enhancer regions (ATAC-seq data by (25)) compared to genome-wide regions lacking ATAC-seq peaks (genomic regions of 2.5 Kb). **(C)** Box plot demonstrating an increase in total GRO-seq read coverage at the putative enhancers (26) in *csr-1* (left) or *drh-3* (right) mutant L3 larva. GRO-seq data (38) were normalized as described in methods, and log2 transformed RPKM was plotted. P-values <2.2*10^−16^ (Wilcoxon test). Cumulative plots confirming the GRO-seq data analysis for *csr-1* (on the left) and *drh-3* (on the right) mutants and demonstrating an increase in GRO-seq reads coverage in both mutants at predicted enhancers (26). Analysis of cumulative distribution of log2-transformed fold changes of GRO-seq RPKM values was performed for genome-wide regions not containing putative enhancers (all chromatin domains) and for combined domains 8, 9 and 10 (26) containing putative enhancer regions. **(D)** GRO-seq reads corresponding to antisense transcription are shown at the region containing a regulatory element detected by ATAC-seq upstream of the *daf-12* gene. An increase in the number of antisense reads is shown in *drh-3* (blue) and *csr-1* (green) mutants compared to wild type (grey).

Importantly, in both *csr-1* and *drh-3* mutants, we found an increase in H3K9me3 at enhancer regions defined by ATAC-seq (25) (Figure 3A). Therefore, it is possible that elevated ncRNA transcription, including antisense transcription, directly or indirectly, leads to chromatin compaction at these regions. Perhaps this mechanism involves dsRNA production and RNAi-induced silencing that depends on non-canonical small RNAs. Notably, the role of nuclear RNAi in suppressing overlapping transcription of convergent genes and read-through antisense transcription had already been described in fission yeast (63,64).

### Rare enhancer small RNAs may guide histone modifications in the absence of secondary siRNAs

To examine the mechanism underlying enhancer silencing, we searched for small RNAs originating from enhancers in published datasets (27,36). Intriguingly, there was larger number of small RNAs mapping to the intronic enhancer of the cyclin E gene in the *csr-1* and *drh-3* mutant datasets (Figure 4A). Notably, the most prominent peak of sense-oriented small RNAs mapped to the annotated tRNA gene present in this intron (Figure 4A). Some additional enhancer small RNAs mapped to known intronic ncRNAs (Figure S2A).

**Figure 4.**
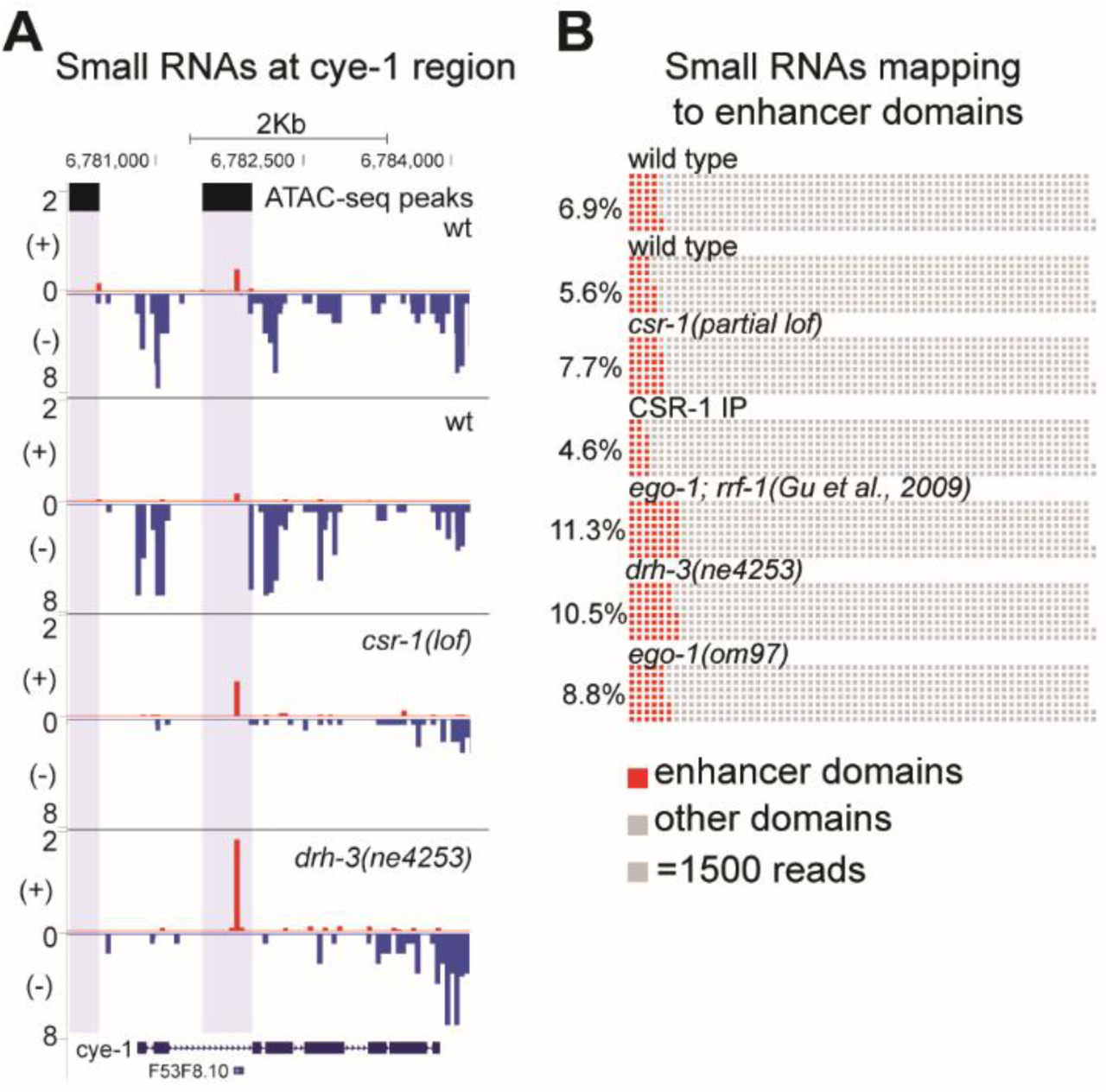
Small RNA increases at enhancer regions in *csr-1* and RdRP mutants compared to wild type. **(A)** Distribution of small RNAs at *cye-1* gene region. Normalization was performed as described in methods. The gene is transcribed from left to right. The reads on the minus strand (in blue) corresponding to 22G-small RNAs are abundant in wild type (two upper tracks), but are depleted in mutants (two lower tracks). On the contrary, there is a sharp peak of small RNAs at the intronic enhancer region (marked with grey), detected by (25), on the plus strand in mutants compared to wild type. ATAC-seq peaks (25) are represented by the black squares with extended grey background. Waffle plots representing small RNA presence at enhancer regions. **(B)** Small RNA read coordinates were intersected with enhancer domain coordinates (26). Percentages of reads were obtained and grouped in blocks with each block representing 1500 reads. Small RNA reads mapped to enhancer regions are represented by red blocks and the percentage is shown on the left.

In addition to the *csr-1* and *drh-3* mutant datasets that are only partially depleted of secondary siRNAs, we have also included into analysis small RNAs cloned from strong RdRP-deficient mutants that lack either one RdPR, EGO-1, required for germline RNAi (65), or both EGO-1 and RRF-1 RdRPs (27).

In wild type animals, about ~7% of small RNA reads map to the enhancer domains defined by (26). Moreover, they are present in the CSR-1 IP samples, i.e. are bound by the CSR-1 Argonaute. Interestingly, 22G-RNA depletion in the RdRP mutants is associated with a notable increase in abundance (to 9-11%) of the “enhancer” fraction of small RNAs (Figure 4B and Figure S2B). It is possible that less abundant small RNAs are cloned more efficiently under these conditions. However, another possibility could be that rare small RNAs may have a better chance to interact with Argonautes lacking their more prominent partners. This could explain an increase in H3K9me3 at enhancers in *csr-1* and *drh-3* mutants (Figures 4B, S3 and 6B) and would suggest that WAGO could be responsible, perhaps in a redundant manner.

### Histone H3K27 methylation is ectopically increased on highly active genes upon *drh-3* loss-of-function

Our H3K27me3 and H3K36me3 ChIP-seq results obtained using wild type worms are consistent with the previous studies (66–68) and demonstrate that H3K36me3 is linked to actively transcribed genes, while H3K27me3 is enriched in regions with low transcription activity (Figure 5A).

**Figure 5.**
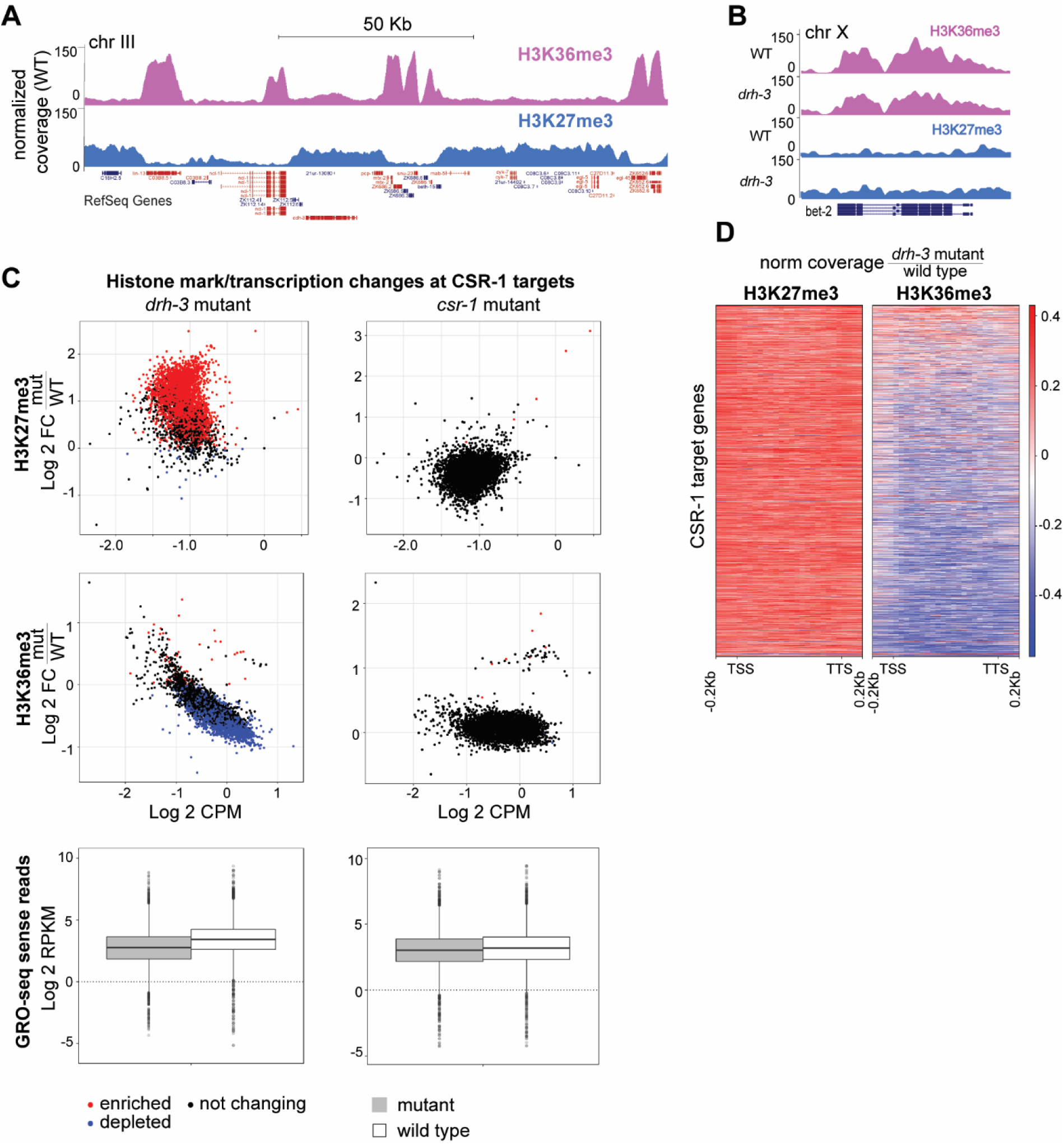
Ectopic H3K27me3 accumulation at highly expressed genes in *drh-3(ne4253)* mutants. **(A)** Open chromatin is associated with H3K36me3 histone modification (pink), which opposes silencing histone mark H3K27me3 (blue). Highly expressed genes (among them CSR-1 targets: *ncl-1*, *lin-13*, *snu-23*) are enriched for H3K36me3 and depleted of H3K27me3. H3K27me3 is abundant at low expressed/silent regions. Highly expressed genes are in red, low expressed genes (expression quantile 1 and 2, see supplementary figure S4A) are in blue. The peaks were generated with MACS2 peak calling algorithm (Galaxy version 2.0.10.2) (Zhang et al., 2008). **(B)** ChIP-seq normalized coverage for H3K36me3 (pink) and H3K27me3 (blue) demonstrating a decrease in H3K36me3, in *drh-3(ne4253)* mutant compared to wild type, and an increase in H3K27me3 modification at the highly expressed CSR-1 target gene *bet-2*. **(C)** Global increase in H3K27me3 and reduction of H3K36me3 at highly expressed CSR-1 target genes in *drh-3(ne4253)* mutant, but not in partially rescued *csr-1(tm892)* mutant (scatterplots), concordant with the reduction of number of GRO-seq reads (38) reflecting the nascent transcript amount (box plots) in mutants compared to wild type. For histone modification analysis, the normalization of both replicas was performed as described in materials and methods. Only the subclass of highly active genes (CSR-1 targets) is represented. The colors reflect the histone mark enrichment (red) and depletion (blue) observed in both biological replicates. GRO-seq data (38) were normalized as described in methods, and log2 transformed RPKM was plotted. P-values: for *drh3(ne4253)* mutant vs wt p-value <2.2*10^−16^, for *csr-1(tm892)* mutant vs wt p-value=1.092*10^−11^(Wilcoxon test). **(D)** Heatmap demonstrating an increase in heterochromatic modification H3K27me3 and a decrease in activating modification H3K36me3 at actively transcribed gene coding regions in the *drh-3(ne4253)* mutant compared to wild type. Each row corresponds to a CSR-1 target gene body between transcription start site (TSS) and transcription termination site (TTS) +/- 200 bp. Different gene length was approximated to be 1000 bp for the representation purpose. The mean value of 2 biological replicates was plotted.

In *drh-3(ne4253)*, transcription reduction of normally highly expressed genes strongly correlates with a reduction in the levels of active histone modification H3K36me3 and a gain of repressive histone mark H3K27me3 (Figure 5B). Many highly expressed mRNAs are known to be complementary to CSR-1-bound small RNAs, i.e. they represent CSR-1 target genes (Figure S4A)(36). Therefore, the reduced expression of CSR-1 target genes in *drh-3(ne4253)* mutant correlates with a global reduction in H3K36me3 accompanied by a gain in H3K27me3 (Figure 5C, left panel and Figure 5D).

In order to confirm that changes in the methylation patterns are not random and linked to transcription alterations, we examined two random sets of genes (4000 and 5000) comparable to the number of CSR-1 target gene population. For this, we first excluded mitochondrial RNA, structural RNA, miRNA and 21U-RNA genes, and then listed all the remaining genes according to their location on the chromosomes and selected every 5^th^ or 4^th^ gene to generate two separate groups (Supplementary Table S3). We did not observe any substantial changes when we compared the mean ChIP-seq coverage for each histone modification at these random sets of genes between *drh-3(ne4253)* and wild type (Figures S4B and C).

At the same time, we observed that transcriptionally silent or less active regions (pseudogenes and tissue-specific genes, represented by a group of spermatogenesis genes), which become more active in *drh-3(ne4253)*, gain H3K36me3 and lose H3K27me3 in the *drh-3(ne4253)* mutant (Figures S4D and E).

Surprisingly, in the partially rescued *csr-1(tm892)* mutant (strain WM193, (36)), we did not observe dramatic changes in H3K36me3 and H3K27me3 (Figure 5C, right panel), even though transcription changes similar to those occurring in *drh-3(ne4253)* take place in this mutant as well (38).

Overall, our results indicate that a loss of “activating” siRNA function leads to ectopic H3K9me3 at CSR-1 target genes, likely dependent on WAGO (Figures 1 and 6A). However, a loss of both “activating” and “silencing” siRNAs results in dramatic redistribution of H3K36me3 and H3K27me3 histone modifications along gene bodies (Figures 5 and 6A). Since WAGO siRNAs are depleted in the *drh-3(ne4253)* mutant, we favor the possibility of additional mechanisms responsible for ectopic deposition of H3K27me3.

**Figure 6.**
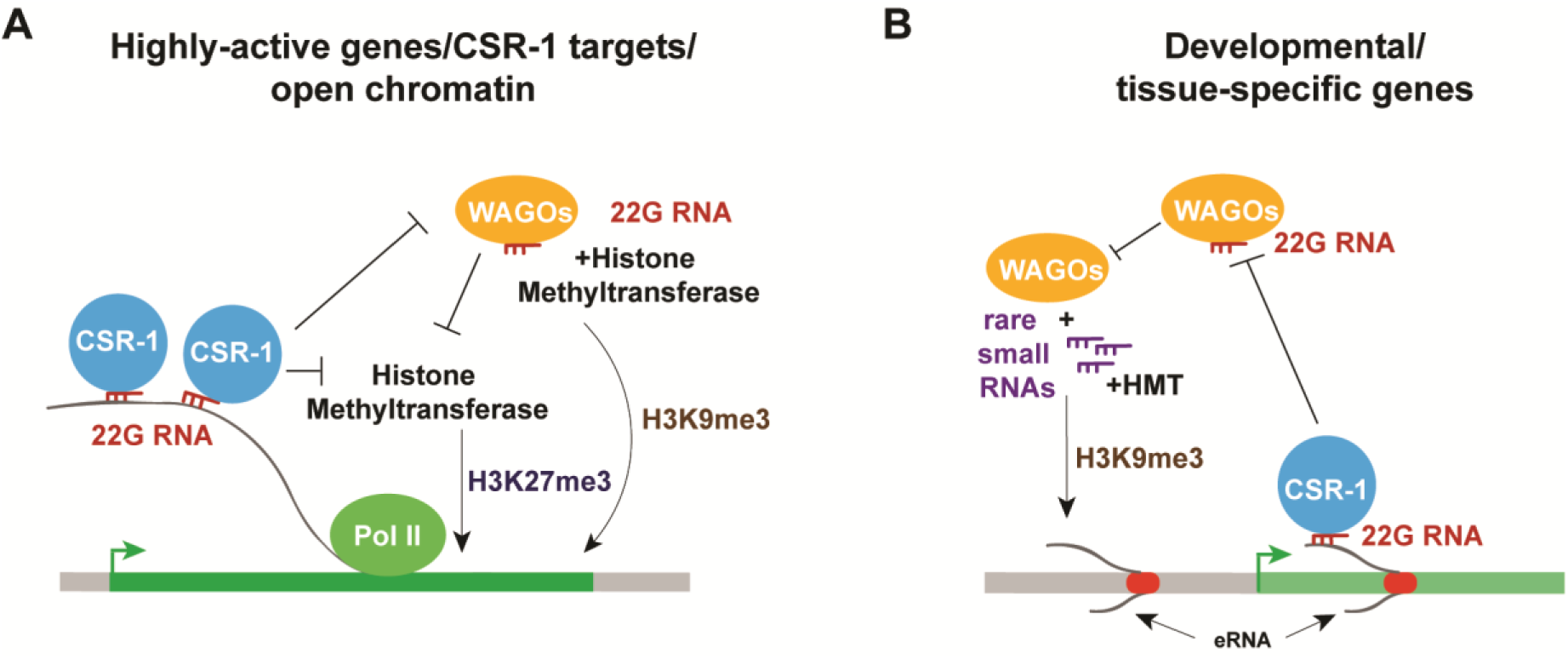
Models describing connections between small RNAs and chromatin at actively transcribed genes (A) and enhancer regions (B). **(A)** 22G-RNAs in complex with CSR-1 Argonaute bind nascent transcripts of highly active genes, thus protecting the latter from WAGO-bound 22G-RNAs guiding H3K9me3 associated with silencing. Either active transcription facilitated by CSR-1 Argonaute, or silencing by WAGO (when CSR-1 is deficient) prevent H3K27me3-modulated silencing through additional mechanisms. **(B)** CSR-1-associated 22G-RNAs bind some enhancer-derived transcripts protecting them from silencing by WAGO. In the absence of either CSR-1 Argonaute or secondary 22G-siRNAs, WAGO may load rare small RNAs that accumulate in these regions. This may lead to WAGO-linked silencing through methylation of H3K9.

## DISCUSSION

Nuclear RNAi in *C. elegans* has been associated with secondary 22G-RNAs generated in the cytoplasm that then move to the nucleus to execute their function. It has been largely studied using exogenous dsRNA for silencing initiation and reporter sequences for nuclear silencing readout (6,8,28–33,69).

Our work links deregulation of the nuclear 22G-RNA pathways to ectopic H3K9 and H3K27 methylation at endogenous genes. Both histone marks can be artificially induced on active genes targeted by exogenous dsRNA (8,31,70), and several H3K9 methyltransferases have been connected to WAGO-induced silencing (30,70–72). However, it was not possible to reliably correlate nuclear Argonautes’ function in inhibiting endogenous genes with H3K9 methylation (34,35). Our results connect an increase in H3K9me3 to reduced transcription of CSR-1 target genes when CSR-1 is limited (Figure 6A). Our data also support earlier models suggesting that under these conditions WAGO may interact with pre-mRNAs normally engaged by CSR-1 (59,60,73). Consistent with this possibility, a deeper analysis of small RNAs present in the WAGO IP samples identified a considerable amount of 22G-RNAs matching typical CSR-1 targets in wild type worms (74), and dramatic misrouting of 22G-RNAs normally bound by CSR-1 to HRDE-1 takes place in the absence of piRNAs (75).

Given the role of DRH-3 (i.e. 22G-RNAs) in maintaining normal levels of H3K9me3 (Figure 1), we were surprised to find an increase in H3K9me3 at enhancer elements in both *csr-1* and *drh-3* mutants. Notably, we found small RNAs matching enhancers in previously published datasets, including CSR-1 IP samples (27,36). A logical explanation of these results is that, in the absence of 22G-RNAs or when CSR-1 is limited, enhancer small RNAs may get loaded onto silencing WAGO that are working together with H3K9 methyltransferases (Figure 6B). It is known that tRNA fragments can be bound by Argonautes (76), and we identify tRNA-derived and ncRNA-derived small RNAs at the intronic enhancers in *C. elegans*. It is possible that they do work with CSR-1 in preventing chromatin compaction at both intragenic and distal enhancer locations.

We would like to emphasize that in this study, as well as in our earlier GRO-seq publication (38), we used a partially rescued *csr-1* mutant strain WM193, (36), which is homozygous viable. Moreover, we used L3 stage worms with the intact germline. These conditions are distinct from those utilized by Desai and colleagues for studying cytoplasmic CSR-1 function in the early embryo (77), and also from those in our report on the role of CSR-1 in histone mRNA maturation, which employed *csr-1(RNAi)* in the sensitized background during post-embryonic development (78).

A surprising finding that should stimulate further inquiry is that the most dramatic histone modification re-distribution involving H3K36me3 and H3K27me3 was observed in the *drh-3(ne4253)* mutant background, where both CSR-1 and WAGO pathways are largely inactive. This suggests that either 22G-RNA-independent silencing mechanisms are involved (Figure 6A), or that residual 22G-RNAs present in *drh-3(ne4253)* play some role (27). The latter possibility may involve WAGO-1, which was connected to small RNA-guided H3K27me3 (70). In either case, it is likely that observed elevation of antisense RNA expression is functionally important. First, it may lead to dsRNA formation and nuclear RNAi initiation through Dicer-dependent siRNA production that may be followed by limited 22G-RNA generation and WAGO-1-guided H3K27me3. Alternatively, nuclear antisense or other ncRNA may participate in promoting H3K27me3 by acting as a scaffold for histone modifying complexes in scenarios similar to those described in other systems (79).

Overall, we uncovered an unexpected complexity in the ways that small RNAs affect chromatin in *C. elegans*. In addition to providing evidence for the competition between the CSR-1 and WAGO pathways in regulation of most active genes in the nucleus, we describe their involvement in shaping enhancer chromatin and also connect nuclear RNAi to regulation of H3K27 methylation on a genome-wide scale.

## Supporting information

## SUPPLEMENTARY DATA

Supplementary Data are available at NAR online.

## ACKNOWLEDGEMENTS

We are grateful to the Mello labs for providing RNAi mutant strains, and we thank Daniel Cifuentes, Dmitry Kretov and members of the BU RNA Club for fruitful discussions.

## CONTRIBUTIONS

E.G. conducted experiments and analyzed data, R.E. analyzed data, Q.M. and N.L. analyzed small RNA data, A.G. supervised the project. E.G. and A.G. wrote the manuscript.

