## Supplementary material for "The interplay between small RNA pathways shapes chromatin landscapes in *C. elegans*"

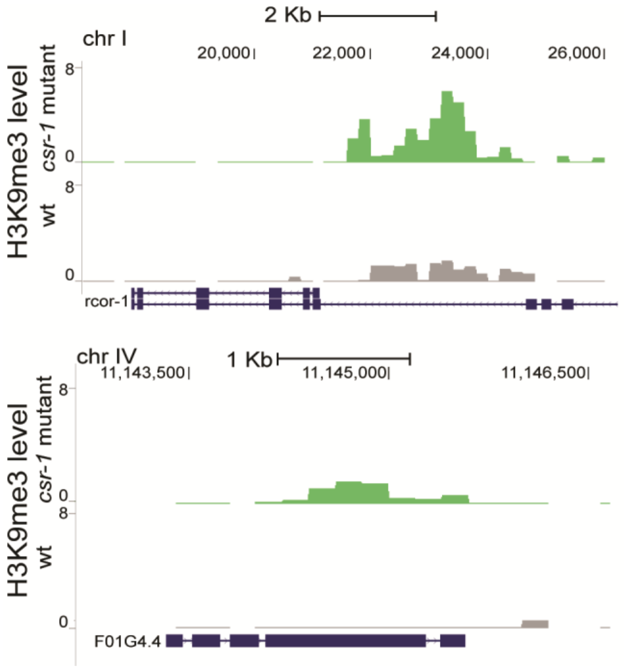


**Supplementary figure S1, related to Figure 1.** Increased H3K9me3 at CSR-1 targets genes in partially loss-of function mutant *csr-1(tm892).* Coverage tracks demonstrating two CSR-1 target genes of different size and exon/intron composition. In wild type (grey), H3K9me3 is very low (upper plot) or is not detected (lower plot), whereas in *csr-1(tm892)* mutant (green), an increase in H3K9me3 is observed. TMM and CPM normalization was performed followed by input reads subtraction for each sample.

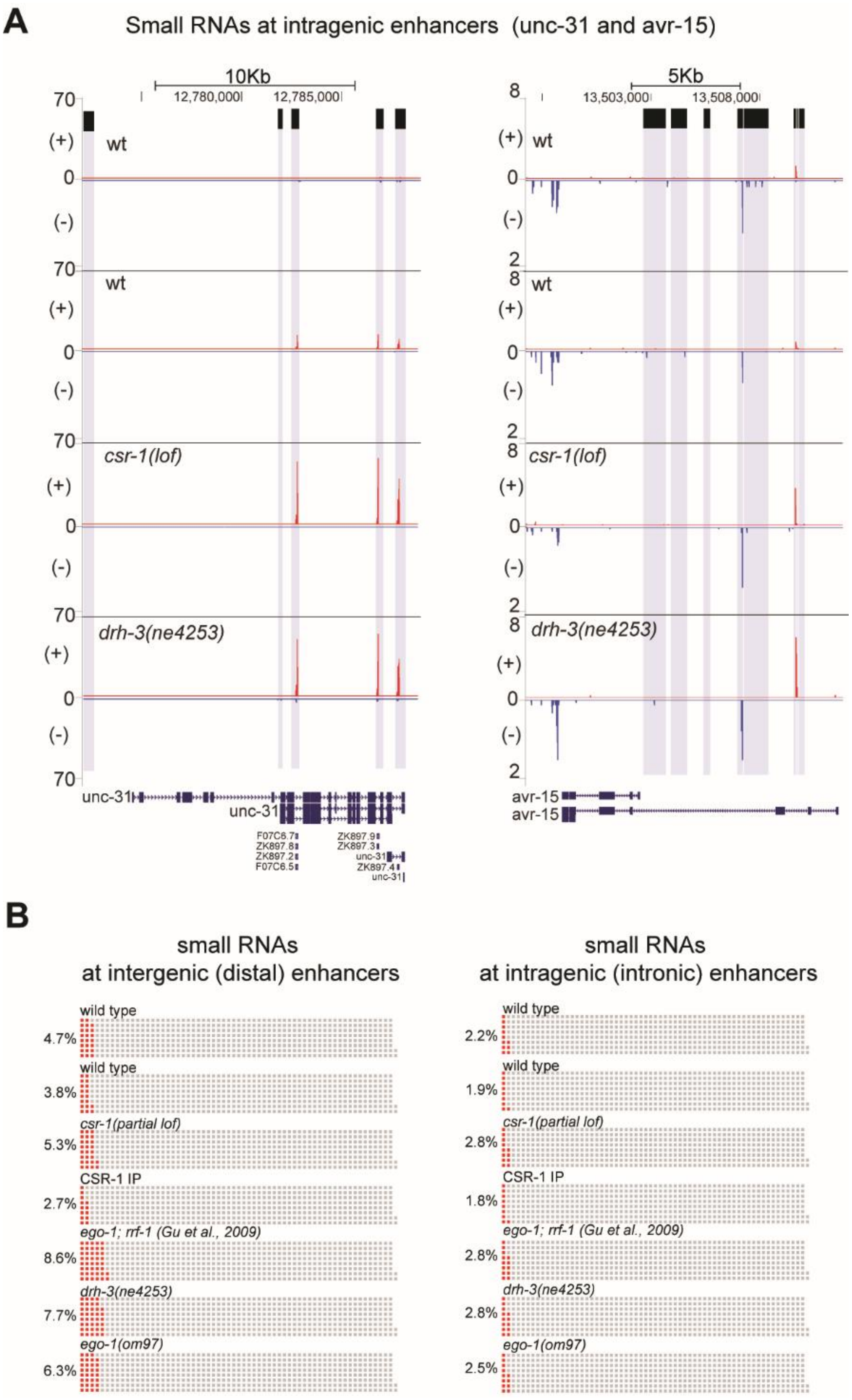

A
Small RNAs at intragenic enhancers (unc-31 and avr-15)
10Kb
12,780,000
12,785,000
5Kb
13,503,000
13,508,000
wt
wt
csr-1(lof)
drh-3(ne4253)
unc-31
avr-15
B
small RNAs at intergenic (distal) enhancers
small RNAs at intragenic (intronic) enhancers
wild type
4.7%
2.2%
wild type
3.8%
1.9%
csr-1(partial lof)
5.3%
2.8%
CSR-1 IP
2.7%
1.8%
ego-1; rrf-1 (Gu et al., 2009)
8.6%
2.8%
drh-3(ne4253)
7.7%
2.8%
ego-1(om97)
6.3%
2.5%

**Supplementary figure S2, related to Figure 4.** Small RNA increases at enhancer regions in *csr-1* and RdRP mutants compared to wild type. **(A)** Distribution of small RNAs at intronic enhancers of *unc-31* (on the left) and *avr-15* (on the right) genes. Normalization was performed as described in methods. The gene *unc-31* is transcribed from left to right; *avr-15* is transcribed from right to left. The reads corresponding to small RNAs at enhancer regions (marked with grey), detected in (25), are more abundant in mutants compared to wild type. ATAC-seq peaks (25) are represented by the black squares with extended grey background. **(B)** Small RNAs map to putative enhancer regions. Waffle plots representing small RNA presence at intergenic and intragenic enhancer regions. Small RNA read coordinates were intersected with enhancer domain coordinates (26). Percentages of reads were obtained and grouped in blocks with each block representing 1500 reads. Small RNA reads mapped to enhancer regions are represented by red blocks and the percentage is shown on the left.

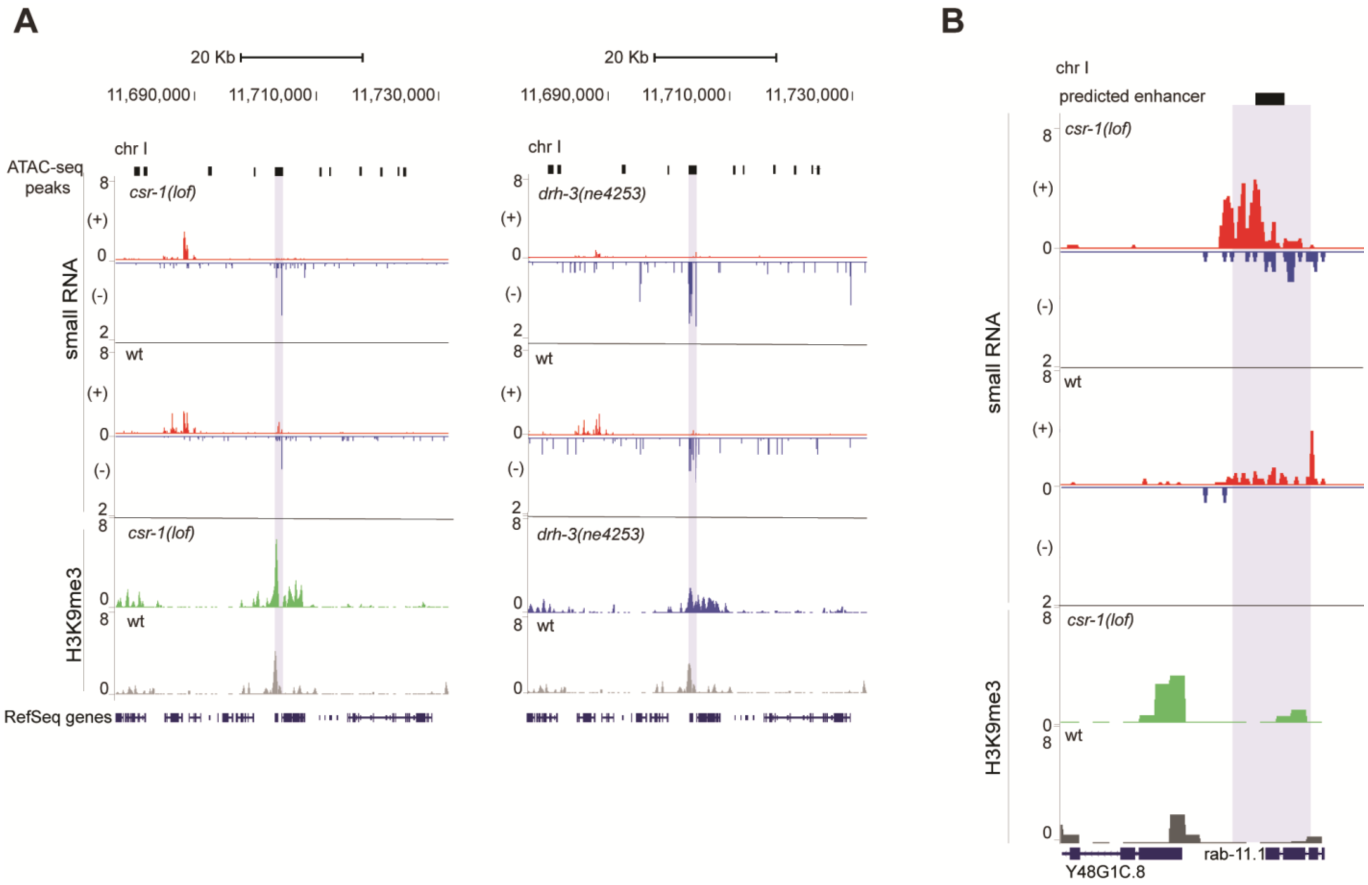


**Supplementary figure S3, related to Figure 4.** Correlation between rare small RNA accumulation around enhancer regions with the increased H3K9me3. **(A)** Distribution of small RNAs at enhancer region on chromosome I in *csr-1(tm892)* mutant (on the left) and *drh-3(ne4253)* (on the right). Normalization was performed as described in methods. The reads corresponding to small RNAs on (-) minus strand (blue) at enhancer region (marked with grey), detected in (25), are more abundant in mutants compared to wild type. Correspondingly, H3K9me3 peak level increases in height and H3K9me3 is spreading around enhancer region in *csr-1(tm892)* mutant (green) and *drh-3(ne4253)* (blue) compared to wild type (grey). ATAC-seq peaks (25) are represented by the black squares with extended grey background. **(B)** Increased abundancy of small RNA around enhancer in *csr-1(tm892)* mutant compared to wild type correlates with the increase in H3K9me3 at this region in *csr-1(tm892)* mutant (green) compared to wild type(grey).

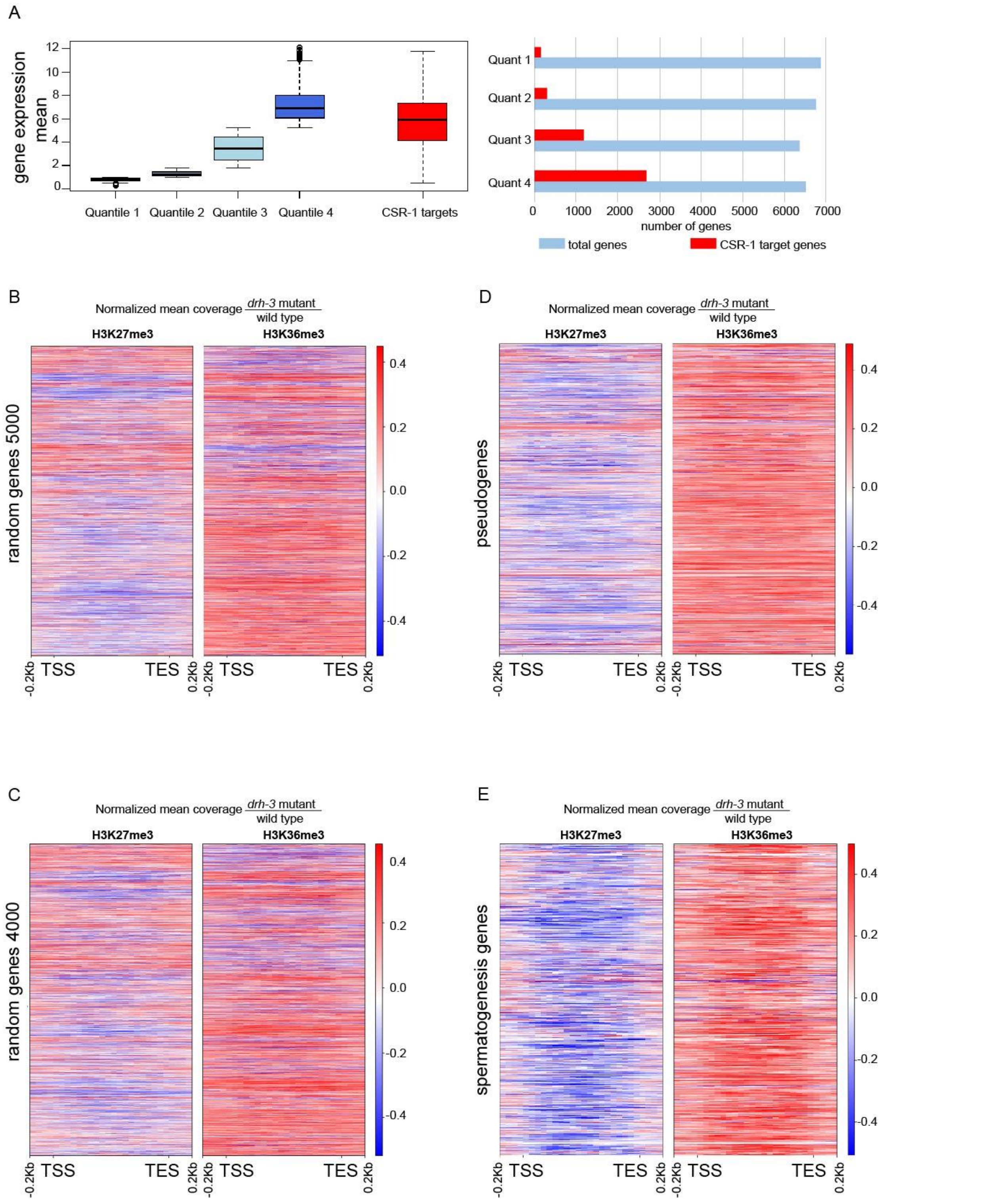


**Supplementary figure S4, related to Figure 5.** Gene expression levels and redistribution of histone modifications in *drh-3* mutant compared to wild type. **(A)** On the left, boxplots showing all

*C. elegans* transcripts (only one isoform of each is represented) divided into four groups corresponding to the level of transcription. The data used for the analysis is obtained from microarray experiment GSE115677. Quartiles were generated using statistical R function “quantile ()”: the genes in quantile 1 represent the lowest level of transcription, those in quantile 4 – the highest. Additionally, the boxplot demonstrates the mean of CSR-1 target gene transcription, which falls in the category between quantiles 3 and 4. To obtain the CSR-1 targets transcription level, CSR-1 dataset (36) was overlapped with the *C. elegans* microarray data (GSE115677). On the rght, the graph demonstrating the number of CSR-1 targets genes(red) compared to the total number of genes(blue) in the quantile. The majority of CSR-1 targets overlap with the genes in quantile 4, i.e. the highly active genes. **(B)** and **(C)** Heatmaps demonstrating the absence of uniform changes in the random sets of genes in *drh-3(ne4253)* compared to wild type. The genome was divided into 100 bp bins and the ratio of normalized bin coverage between mutant and wild type was plotted on a heatmap. Each row corresponds to gene coding regions between transcription start site (TSS) and transcription termination site (TTS) +/- 200 bp. Different gene length was approximated to be 1000 bp for the representation purpose. The mean value of 2 biological replicates is shown. The sets of genes can be found in the Supplementary Table 3). **(D)** Heatmap demonstrating a decrease in silencing modification H3K27me3 and an increase in activating modification H3K36me3 at pseudogenes in *drh-3(ne4253)* compared to wild type. The genome was divided into 100 bp bins and the ratio of normalized bin coverage between mutant and wild type was plotted on a heatmap. Each row corresponds to pseudogene coding regions between transcription start site (TSS) and transcription termination site (TTS) +/- 200 bp. Different gene length was approximated to be 1000 bp for the representation purpose. The mean value of 2 biological replicates is shown. The list of pseudogenes was obtained from Ensembl database and can be found in the Supplementary Table 3. The increase of pseudogene transcription in *drh-3(ne4253)* is previously demonstrated in the work of W. Gu and co-authors (27). **(E)** Same as on (D) for tissue-specific genes represented by spermatogenesis genes. The list of spermatogenesis genes can be found in the Supplemental Table 3. The increase of sense-oriented transcription in spermatogenesis genes *in drh-3(ne4253)* was shown previously (38).
